# Antibiotic degradation by commensal microbes shields pathogens

**DOI:** 10.1101/870931

**Authors:** Mergim Gjonbalaj, James W. Keith, Mytrang Do, Tobias M. Hohl, Eric G. Pamer, Simone Becattini

## Abstract

The complex bacterial populations that constitute the gut microbiota can harbor antibiotic-resistance genes (ARGs), including those encoding for β-lactamase enzymes (BLA), which degrade commonly prescribed antibiotics such as ampicillin. While it is known that ARGs can be transferred between bacterial species, with dramatic public health implications, whether expression of such genes by harmless commensal bacterial species shields antibiotic-sensitive pathogens *in trans* by destroying antibiotics in the intestinal lumen is unknown. To address this question, we colonized GF mice with a model intestinal commensal strain of *E. coli* that produces either functional or defective BLA. Mice were subsequently infected with *Listeria monocytogenes* or *Clostridioides difficile* followed by treatment with oral ampicillin. Production of functional BLA by commensal *E. coli* markedly reduced clearance of these pathogens and enhanced systemic dissemination during ampicillin treatment. Pathogen resistance was independent of ARG acquisition via horizontal gene transfer but instead relied on antibiotic degradation in the intestinal lumen by BLA. We conclude that commensal bacteria that have acquired ARGs can mediate shielding of pathogens from the bactericidal effects of antibiotics.

## Importance

The wide use of antibiotics in human populations and in livestock has led to increasing prevalence of pathogenic and commensal bacterial species that harbor antibiotic resistance genes (ARGs), such as those encoding for ampicillin-degrading β-lactamases. We investigated whether harmless autochthonous bacteria might degrade orally administered antibiotics, thereby impairing their ability to combat intestinal pathogens. Here we report that antibiotic degradation by a resident intestinal strain of *E. coli* reduces the effectiveness of oral ampicillin against two intestinal pathogens, *L. monocytogenes and C. difficile*, resulting in increased intestinal and systemic bacterial burden. We demonstrate that expression of ARGs by non-pathogenic members of the gut microbiota shields antibiotic-sensitive pathogens and enhances their expansion and dissemination.

## Observation

Antibiotic administration has markedly reduced the morbidity and mortality associated with bacterial infections in the pre-antibiotic era. Increasing antibiotic-resistance in pathogenic microbes, mediated in part by acquired genes that encode antibiotic-degrading enzymes, represents a major threat to human health (1).

The gut microbiota contains trillions of commensal bacteria that can also harbor antibiotic resistance genes (ARGs) (2). Notably, antibiotic exposure can increase ARG gene representation and expression by the gut microbiota (3). Horizontal ARG transfer represents a mechanism by which drug-sensitive microbes can acquire resistance, e.g. by acquisition of genes encoding antibiotic-degrading hydrolases (4, 5). Thus, it is possible that commensal bacterial species transfer ARGs to intestinal pathogens upon antibiotic exposure in the gut lumen. However, another possibility is that production of antibiotic-degrading enzymes by the resident microbiota protects otherwise drug-sensitive pathogens *in trans*, thereby facilitating their replication and spread in the host.

To test this hypothesis in a controlled system, we reconstituted germ-free (GF) mice with an *E. coli* strain, utilized here as a model commensal, that expresses either a WT form of β-lactamase (TEM-1) or an inactive point mutant (hereby referred to as *WT* BLA or *mut* BLA, respectively) (Figure 1A) (6). This approach yielded cohorts of mice that, with the exception of one codon, harbor identical genomes, thus excluding differences in microbiota functions (e.g., immune activation, colonization resistance, etc.) that are not related to the β-lactam degradation.

**Figure 1.**
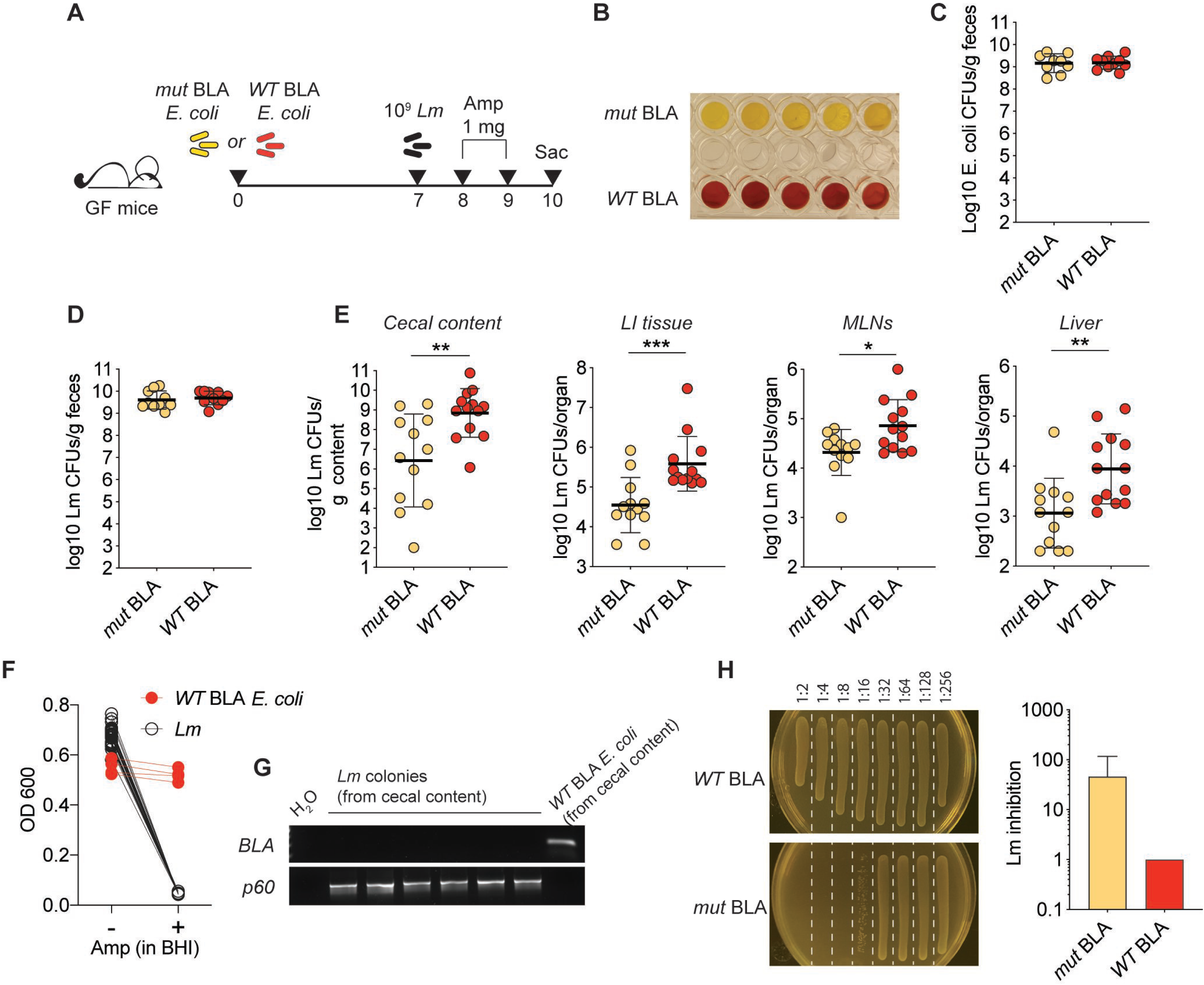
β-lactamase production by a model commensal curtails the efficacy of ampicillin against *Listeria monocytogenes*. **A)** Schematic representation of the experimental design. **B)** Nitrocefin assay performed on resuspended fecal pellets obtained from the depicted groups of mice. Each well represents a different mouse, one representative experiment of three shown. **C)** Reconstitution levels for the depicted *E. coli* strains as measured by plating fecal pellets on day 7 post-reconstitution (day of infection) onto selective plates (n=8-10, data pooled from 2 independent experiments, shown are individual data points and geometric mean). **D)** Luminal *Lm* burden in the depicted mice 1 day post infection, measured by plating fecal pellets onto selective plates. **E)** *Lm* burden in the depicted compartments at day 3 post infection (D,E: n=12, data pooled from 3 independent experiments, shown are individual data points and geometric mean. Mann-Whitney test: *=p<0.05, **=p<0.01, ***=p<0.001). **F)** Individual *Lm* colonies (28) or *WT* BLA *E. coli* colonies (4) from 4-5 different mice were inoculated into BHI +/- ampicillin. OD was measured after o.n. culture. **G)** Colonies utilized for the experiment depicted in F) were also subjected to PCR with primers specific for the TME-1 β-lactamase gene or p60 (*Lm* positive control). Shown are results for 6 *Lm* colonies and 1 *E. coli* colony; identical results were obtained for all tested colonies. H) The cecal content of WT mice reconstituted with either *WT* or *mut* BLA *E. coli* and administered ampicillin in drinking water for two days was serially diluted and inoculated with *Lm. Lm* growth was assessed after o.n. culture by measurement of OD and direct plating (one representative plate shown on the left). Plotted values correspond to the first dilution allowing for detectable *Lm* growth (with 1 indicating Lm growth at all dilutions) (n=3, shown is mean ± SD). Similar results were obtained utilizing antibiotic-treated, *E. coli*–reconstituted animals (see Supplementary Figure 1).

Although *WT* BLA and *mut* BLA *E. coli* reached identical luminal bacterial densities in reconstituted mice, a colorimetric assay confirmed that only the intestinal content of mice reconstituted with *WT* BLA *E. coli* retained the capacity to hydrolyze β-lactams (Figure 1 B, C). One week after reconstitution, mice were orally infected with the foodborne pathogen *Listeria monocytogenes* 10403s (*Lm*). *Lm* is highly sensitive to β-lactam antibiotics and can expand in the gut lumen of mice that lack colonization resistance (7). Mice were then administered ampicillin on day +1 and +2 post *Lm* infection and sacrificed on day +3. As expected, *Lm* reached identical densities in the intestines of *WT* BLA or *mut* BLA *E. coli* reconstituted mice on day +1, indicating that the 2 *E. coli* strains did not differ in their inability to provide colonization resistance against *Lm*. However, we found significantly higher *Lm* burden in multiple organs in mice harboring *WT* BLA *E. coli* on day +3, consistent with the notion that β-lactamase-dependent ampicillin degradation shielded *Lm* from the therapeutic antibiotic’s action. To exclude the possibility that *Lm* might have acquired resistance to ampicillin via horizontal gene transfer, we inoculated single *Lm* colonies recovered from the cecal content of *WT* BLA *E. coli-*reconstituted, *Lm*-infected mice, into liquid medium either in the presence or absence of ampicillin. Notably, none of the inoculated *Lm* colonies grew in the presence of ampicillin, in contrast to *WT* BLA-expressing *E. coli* colonies recovered from the same mice (Figure 1F). Furthermore, none of the *Lm* colonies tested positive for the presence of the β-lactamase gene, which was uniformly detected in colonies of *WT* BLA *E. coli* by PCR (Figure 1G).

To confirm that the increased *Lm* burden observed above was due to antibiotic degradation by resident *E. coli*, we collected the cecal contents of mice reconstituted with either *WT* BLA or *mut* BLA *E. coli* and treated with ampicillin in the drinking water for two consecutive days to allow for luminal accumulation of the antibiotic. Inoculation of *Lm* into serial dilutions of the cecal content supernatants revealed that the cecal contents from mice reconstituted with *mut* BLA *E. coli* had a higher inhibitory capacity compared to cecal contents recovered from mice reconstituted with *WT* BLA *E. coli* (Figure 1G and Supplementary Figure 1). Since the presence of active β-lactamase was the only *bona fide* difference between the cecal contents of the two cohorts of mice, we conclude that the microbiota-encoded enzymatic activity curtailed the efficacy of ampicillin treatment against *Lm*.

To expand our observations beyond the *Listeria* model, and to assess whether commensal-mediated antibiotic degradation may represent a mechanism that is relevant to other infectious agents, we adapted our experimental strategy to an established *C. difficile* infection model (8) (Figure 2A), an important intestinal pathogen that is also sensitive to ampicillin (Supplementary Figure 2). Of note, this model allowed us to investigate the relevance of our findings in a setting where expansion of an antibiotic-resistant microbe takes place following antibiotic-mediated depletion of the intestinal microbiota, a common occurrence in hospitalized patients (9). Similar to the results obtained with *Lm*, we observed indistinguishable levels of expansion for both *E. coli* and *C. difficile* on day +1 after reconstitution or infection, respectively, in all groups of mice (Figure 2B, C). In agreement with our previous findings, the *C. difficile* burden was significantly reduced by ampicillin treatment in mice reconstituted with *mut* BLA *E. coli*, but not in mice reconstituted with *WT* BLA *E. coli* (Figure 2D). Direct comparison of the ampicillin-treated mice confirmed a significantly higher burden in mice whose intestinal flora had the capacity to hydrolyze β-lactams (Figure 2D).

**Figure 2.**
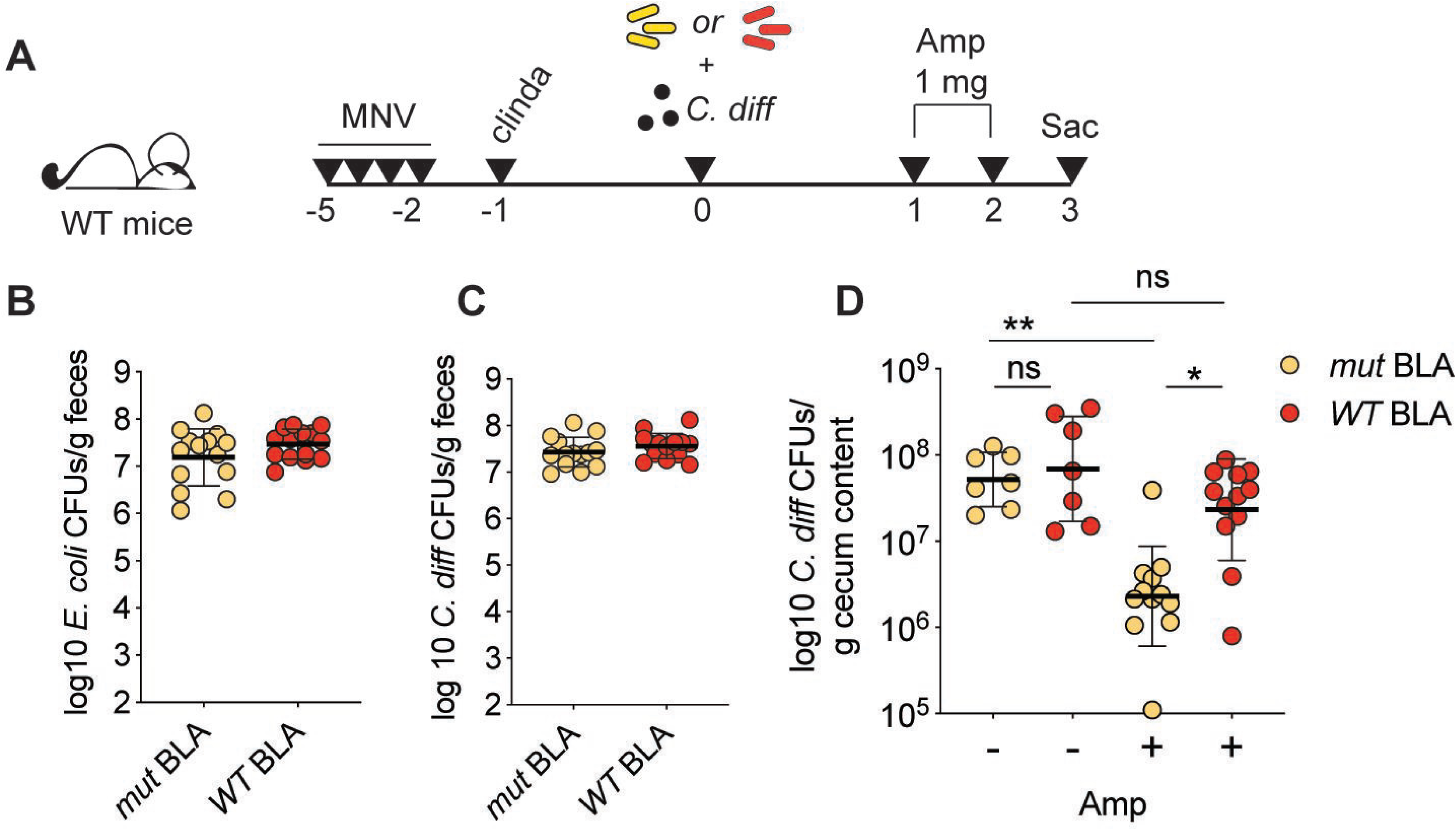
Endogenous antibiotic degradation impacts treatment of *C. difficile* infection. **A)** Schematic representation of the experimental design. **B)** Reconstitution levels for mice reconstituted with either *WT* or *mut* BLA *E. coli* (as depicted in A), one day post oral gavage as assessed by selective plating of fecal pellets (n=14, data pooled from 2 independent experiments, shown are individual data points and geometric mean). **C)** *C. difficile* burden in mice treated as depicted in A), one day post oral gavage, as assessed by selective plating of fecal pellets (n=14, data pooled from 2 independent experiments, shown are individual data points and geometric mean). **D)** *C. difficile* burden in the cecal content of mice treated as depicted in A) at day 3 post infection (n=7 for controls, n=12 for amp-treated, data pooled from 3 independent experiments, shown are individual data points and geometric mean; Kruskal-Wallis test with multiple comparisons: *=p<0.05, **=p<0.01).

These findings suggest that ARGs expressed by commensal bacteria can shape the chemical niche of the intestine and confer an apparent antibiotic resistant phenotype to pathogens in trans, without direct acquisition of ARGs by the pathogenic microbe. We refer to this activity as *commensal-mediated pathogen shielding*. Using two different infection models, we show that production of β-lactamases, a prototypical antibiotic resistance factor, by resident intestinal microbes can significantly reduce the effectiveness of ampicillin treatment, thereby generating a safe environment in which otherwise sensitive pathogens are shielded from this drug. Importantly, previous studies in healthy volunteers demonstrated that upon treatment with cephalosporins, subjects harboring BLA-producing commensal strains, unlike BLA-negative subjects, had undetectable concentrations of the drug in the feces and maintained a rich microbiota, providing evidence that BLA concentrations sufficient to inactivate antibiotics are commonly achieved in humans (10, 11).

Whether or not ARGs enrichment within the gut microbiota is detrimental to host health is a complex question, and the answer is likely to be context-dependent.

For instance, oral administration of recombinant beta lactamase or BLA-producing bacteria was shown to preserve the integrity of the microbiota following parenteral administration of beta-lactam antibiotics in animal models, without affecting drug concentration in the serum (12-17). These approaches were shown to be advantageous in that they preserved colonization resistance against pathogens (12-17).

On the other hand, early work (reviewed in (18)) revealed that beta-lactamase-producing, non-pathogenic bacteria, can hinder the efficacy of penicillins *in vitro* and *in vivo*, using models of subcutaneous and tonsil infection. Clinical data also suggested that the presence of one beta-lactamase-producing bacterial strain at the site of infection could enhance persistence of a pathogen upon antibiotic treatment (19). In these settings, members of the Bacteroides genus, among the most highly represented genera in the human intestine(20), were also identified as BLA-carriers.

Consistent with these observations, our laboratory recently showed that a few bacterial strains, out of the dozens composing the microbiota of a mouse colony treated with ampicillin for over 8 years, had the capacity to hydrolyze ampicillin, while the other bacterial strains, in isolation, remained sensitive to ampicillin and thus were protected *in trans* by a minor subset of the microbiota (21)(see Figure 4A in (22)).

In conclusion, we propose that *commensal-mediated pathogen shielding* can impair the effectiveness of some antibiotic treatments during infection. While pharmacokinetic studies have generally focused on antibiotic absorption, distribution, enzymatic modification, protein binding and biliary/renal clearance, the role of microbiota-mediated antibiotic degradation in the gut lumen and its potential for dramatically impacting responses to antibiotic treatment has received less attention. Our findings extend the recently uncovered broad capacity of the gut microbiota to metabolize drugs, affecting their efficacy (23, 24). Within this model, antibiotics represent an additional class of xenobiotics that commensals can metabolize.

Our study suggests that presence or absence of commensal bacterial strains that inactivate beta-lactam antibiotics is likely to impact clinical responses to antibiotic treatment, possibly contributing to inter-individual variability in therapy outcomes. Furthermore, the occurrence of *pathogen shielding* might be a relevant element to consider in the engineering of probiotic bacterial strains to be employed in clinical practice.

## Methods

### Mouse Husbandry

All experiments using wild-type mice were performed with C57BL/6J female mice that were 6–8 weeks old; mice were purchased from Jackson Laboratories. Germ-free (GF) mice were bred in-house in germ-free isolators. Following reconstitution mice were housed in sterile, autoclaved cages with irradiated food and acidified, autoclaved water. All animals were maintained in a specific-pathogen-free facility at Memorial Sloan Kettering Cancer Center Animal Resource Center. Experiments were performed in compliance with Memorial Sloan-Kettering Cancer Center institutional guidelines and approved by the institution’s Institutional Animal Care and Use Committee.

### Generation of *E. coli* strains

Plasmids encoding for WT TEM-1 βlactamase (pDIMC8-TEM1) or mutated TEM-1 βlactamase (pDIMC8-TEM1 W208G) were extracted from the RH06 and RH09 *E. coli* strains, published elsewhere (6), gel-purified and utilized for transformation of Stellar competent cells (Takara Bio) according to manufacturer’s instructions. The resulting strains were utilized for experiments throughout this study. Of note, the plasmids conferred resistance to chloramphenicol, and while expression of the TEM-1 gene was placed under the regulation of a tac promoter, we did not induced it by IPTG treatment, but rather exclusively relied on leaky transcription of the gene, to produce more physiologically-relevant conditions.

### Antibiotic treatment, reconstitution and infections

GF mice were gavaged with either of two strains of *E. coli*, encoding for a functional or a point-mutated version of TEM-1 β-lactamase, respectively. 1 week post reconstitution mice were gavaged with 10^9^ CFUs of *L. monocytogenes* (*Lm*) strain 10403s and administered 1 mg of ampicillin (Fisher) by oral gavage daily for 2 consecutive days. Animals were euthanized at day 3 post infection. Reconstitution of WT mice with E. coli strains for in vitro experiments involving dilution of cecal content, mice were treated for 3 days with metronidazole and vancomycin in drinking water (0.5 g/l), left on regular water for 1 day, and then gavaged with the appropriate *E. coli* strain. 1 week post reconstitution mice were treated with ampicillin in drinking water (0.5 g/l) for two days prior to being euthanized.

For *C. difficile* infection experiments, WT C57Bl/6 mice were administered a combination of metronidazole, neomycin and vancomycin (0.25 g/l each) in drinking water for 3 days, and 24h post antibiotic regimen cessation were injected i.p. with clindamycin (200 μg). On the following day mice were reconstituted with either *WT* or *mut* BLA *E. coli* (5×10^4^ CFUs) and 200-500 spores of *C. difficile* strain VPI10463 (ATCC #43255).

### CFUs enumeration and Selective plating

*L. monoctogenes* was identified through plating of serial dilutions of homogenized organs (prepared as described elsewhere (7)) or fecal material (resuspended 100 mg/ml in PBS) ontoBHI plates supplemented with streptomycin (100 μg/ml) and nalidixic acid (50 μg/ml).

*E. coli* CFUs were enumerated following plating of serial dilution of fecal material onto LB plates supplemented with chloramphenicol (50 μg/ml). *E. coli* CFU numbers obtained from plating of ex-GF mice at day of infection onto LB plates (not supplemented with antibiotics) yielded identical numbers, indicating that plasmids carrying CM resistance cassette as well as the *WT/mut* BLA gene were maintained even in the absence of any selective pressure.

For detection of *C. difficile*, fecal pellets or cecal content were resuspended in deoxygenated phosphate-buffed saline (PBS), and ten-fold dilutions were plated on BHI agar supplemented with yeast extract, taurocholate, L-cysteine, cycloserine and cefoxitin at 37°C in an anaerobic chamber (Coylabs) overnight.

### *Lm* culture in cecal content

Cecal contents were recovered from *E. coli* reconstituted WT or GF animals, resuspended in PBS at 300 mg/ml (WT) and spun down at 3000 rpm for 10’. Serial 1:2 dilutions of the resulting supernatant were generated using PBS and 100 μl of each dilution were plated in replicate in flat bottom 96 well plates. An equal volume of BHI medium supplemented with streptomycin (200 μg/ml) and nalidixic acid (100 μg/ml) acid (to prevent growth of residual *E. coli*) containing 100-1000 CFUs *Lm* 10403s, was added on top. *Lm* for this assay was prepared by re-inoculating an overnight culture in liquid BHI at 37°C on shaker, until logarithmic phase of growth was reached (OD=0.1-0.4). After an overnight incubation at 37°C, the plate was assayed by OD 600 reading and individual dilutions plated onto BHI-Strep-NA plates for assessment of *Lm* growth. Normalized inhibition index was calculated as 1/first dilution allowing for *Lm* growth, with the initial dilution being 1:2 to take into account the addition of a volume of BHI equivalent to that of the medium. For example, if the first dilution where *Lm* was detected was 1:16, the resulting inhibition index would be 16. Within each experiment samples were then normalized to the baseline, obtained by averaging the values obtained in the control group, represented by mice reconstituted with *mut* BLA *E. coli*.

### PCR

PCR for was carried out using the following primers: β-lactamase (fw: 5’-GCTATGTGGCGCGGTATTAT-3’; rev: 5’-AAGTAAGTTGGCCGCAGTGT-3’, product: 191 bp); p60 (fw: 5’-GCGCAACAAACTGAAGCAAAGGATGC-3’; rev: 5’- CTCGCGTTACCAGGCAAATAGATGGACG-3’, product: 1300 BP), using the SapphireAMp Fast PCR master mix (Takara Bio) and the following conditions: 94°C × 1’, 30 × (98°C ×; 5’’, 58°C × 5’’, 72°C × 15’’).

## Acknowledgments

We thank Ying Taur and Peter McKenney for critical discussion of the manuscript. RH06 and RH09 *E. coli* strains carrying the pDIMC8-TEM1 and pDIMC8-TEM1 W208G plasmids were a kind gift of Prof. Marc Ostermeier (Johns Hopkins University). This work was supported by NIH grant P30 CA008748 (to MSKCC), Burroughs Wellcome Fund Investigator in the Pathogenesis of Infectious Disease Awards (T.M.H.).

S.B. was supported by an Early Postdoc Mobility Fellowship from the Swiss National Science Foundation (P2EZP3_159083) and an Irvington Fellowship from the Cancer Research Institute (49679).

---

**Supplementary Figure 1.**
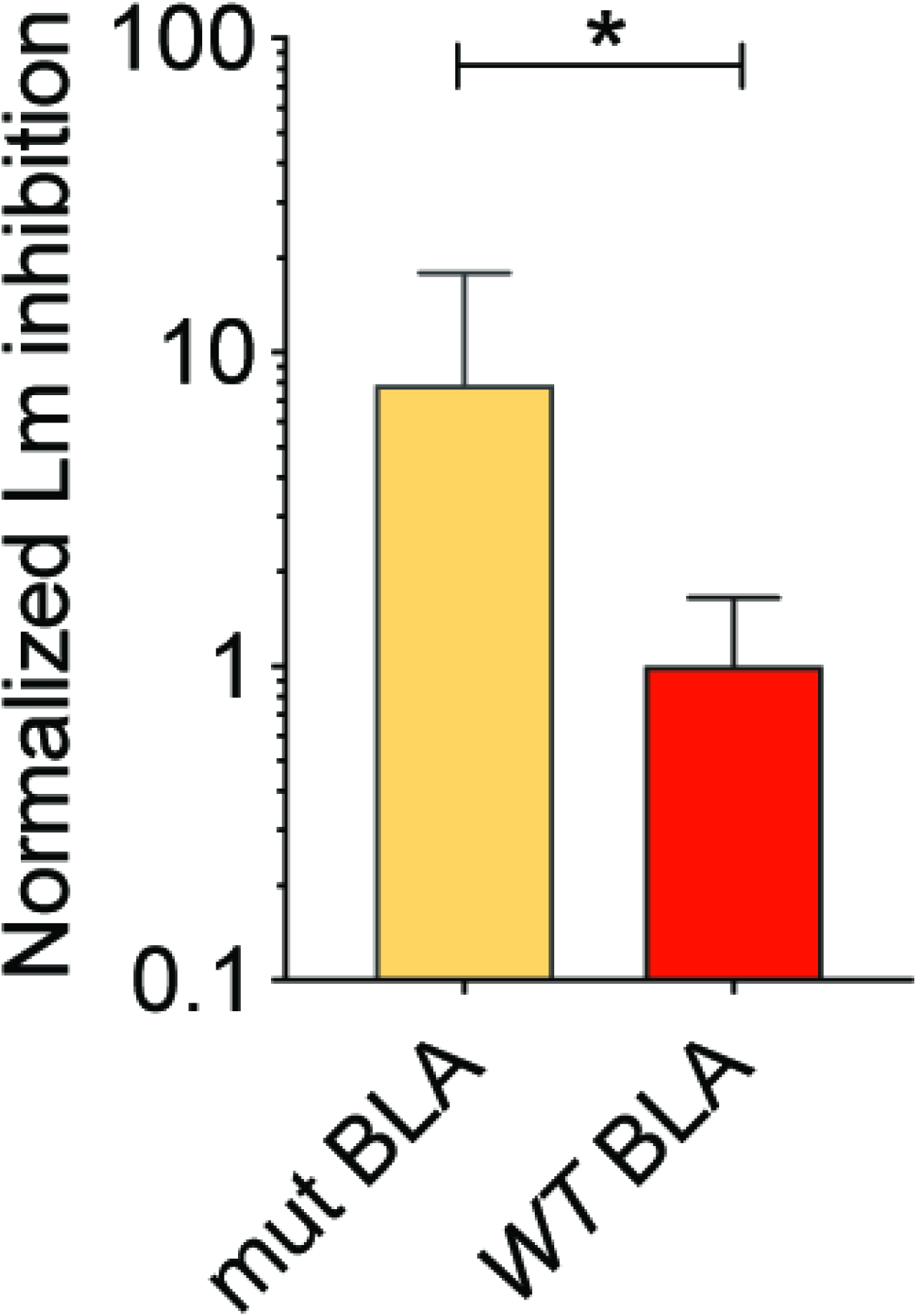
Confirmation of differential anti-listerial activity of cecal contents in *WT* BLA vs *mut* BLA *E. coli*-reconstituted WT mice. C57Bl/6 mice were treated with a combination of metronidazole, neomycin and vancomycin for 3 days and reconstituted with either *WT* or *mut* BLA *E. coli*. 7 days following reconstitution mice were administered ampicillin in drinking water for 2 days, prior to sacrifice. *Lm* was cultured in serial dilutions of the cecal contents, and growth was assessed after o.n. incubation. To account for variability across experiments, *Lm* inhibition calculated as 1/first cecal dilution allowing *Lm* growth was normalized so that the average value for *WT* BLA ceca would be equal to 1 in each experiment (n=9-10 mice from 3 different experiments, shown is mean ± SD, Mann-Whitney test: *=p<0.05).

**Supplementary Figure 2.**
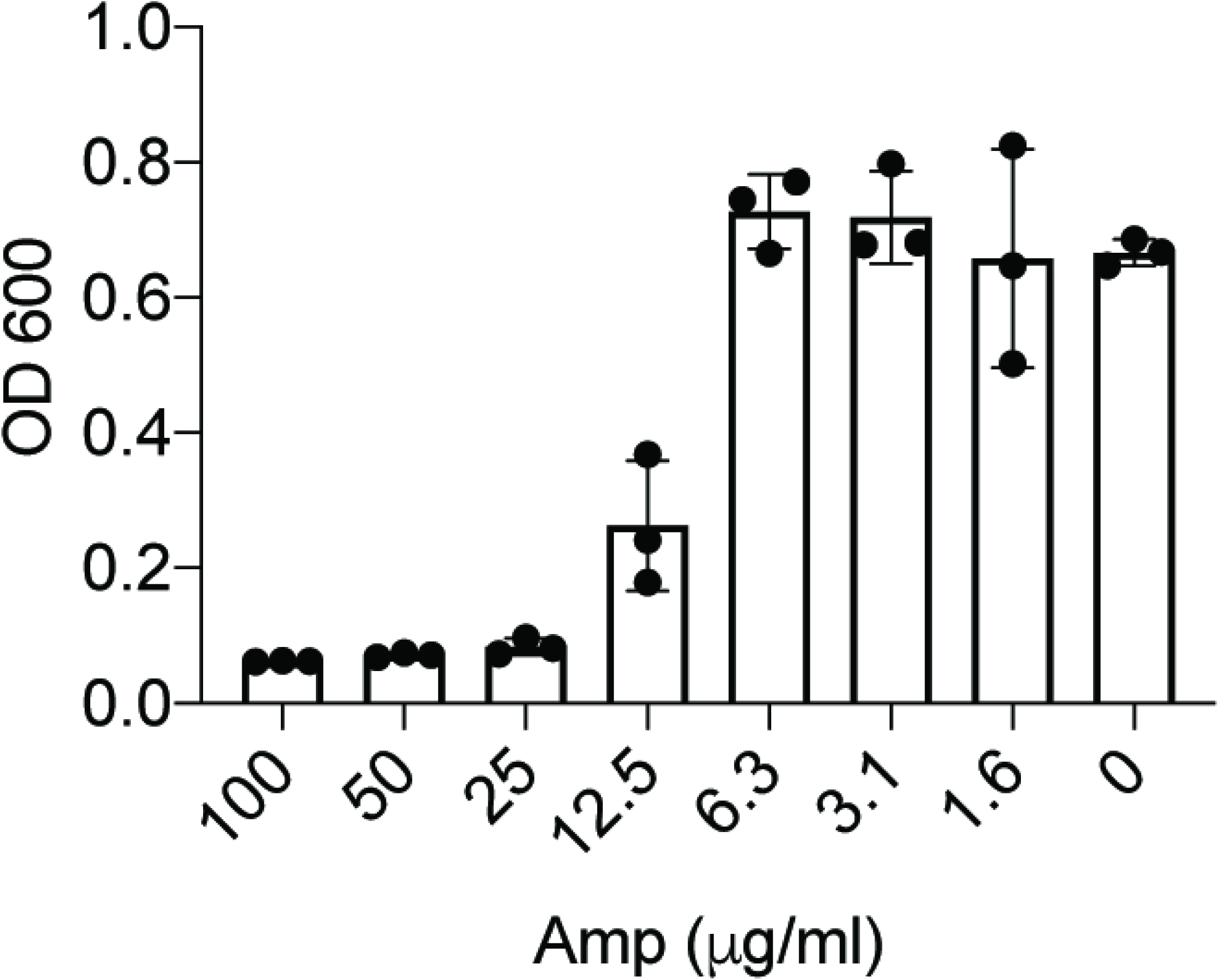
Ampicillin sensitivity of *C. difficile*. *C. difficile* was grown to stationaty phase and inoculated in medium with different concentrations of ampicillin. Shown is OD600 measured after o.n. incubation at 37°C in anaerobic conditions (n=3 technical replicates).

